# microRNA-544a as a new modulator of the Wnt-signalling network in the articular cartilage and osteoarthritis

**DOI:** 10.1101/2024.11.28.625937

**Authors:** Cintia Scucuglia Heluany, Nicholas J Day, Keemo Delos Santos, Anna De Palma, Tracey E Swingler, David Sochart, Ian M Clark, Mohit Kapoor, Giovanna Nalesso

## Abstract

**Objective:** Determining the effect of microRNA-544a (miR-544a) in articular chondrocytes isolated from patients affected by osteoarthritis (OA) and its role in the modulation of the Wnt signalling.

**Methods:** Articular chondrocytes were isolated from patients undergoing joint replacement because of OA. Expression levels of miR-544a were measured by PCR and by in situ hybridization. Putative targets of miR-544a were confirmed by reporter assay and by qPCR in cells stimulated with a miR-544a mimic. The effect of miR-544a on chondrocyte metabolism was monitored by qPCR for phenotypic markers, protein expression levels of aggrecan neoepitopes/MMP-13 and modulation of alcian blue content in micromass cultures, upon stimulation with a miR-544a mimic. The expression levels of MMP-13 and Aggrecan neoepitopes in response to miR-544a stimulation was also measured in co-stimulation with Xav-939 and KN93, which are respectively β-catenin and CaMKII inhibitors.

**Results:** Our results suggest that miR-544a enhances the activation of the Wnt-signalling in the articular chondrocytes, by downregulating the expression of components of the Wnt/β-catenin destruction complex. The expression of miR-544a is higher in chondrocytes isolated from damaged areas of the articular cartilage removed from OA patients, and can be upregulated by pro-inflammatory and pro-fibrotic cytokines. miR-544a exerts a pro-catabolic effect of articular chondrocytes, which is rescued both by the inhibition of the Wnt/β-catenin and Wnt/CaMKII signalling pathways.

**Conclusion:** our results point to miR-544a as a new, important modulator of the Wnt signalling network within the articular cartilage suggesting a key role for microRNAs in regulating how the multiple branches of the network and their interaction modulate cartilage homeostasis.

## Introduction

Osteoarthritis (OA) is a chronic musculoskeletal condition affecting 7% of the global population, with a higher prevalence in females (1). Affected patients can experience severe pain: mobility can be severely impaired, significantly impacting people’s life quality. Progressive degeneration of the articular cartilage, remodelling of the subchondral bone and synovial inflammation are all considered hallmarks of the disease (2).

OA pathogenesis is complex as the disease can be caused by a multiplicity of factors, such as genetic predisposition, history of trauma, obesity, which can contribute to the development of the disease, often in combination, making patient stratification difficult and resulting in a high variability of response to treatment both in pre-clinical and clinical settings.

No disease modifying drug is currently approved for the treatment of OA and surgical joint replacement is offered to patients most severely affected.

Deregulation of several signalling cascades has been associated with the onset and progression of OA. Deregulation of the Wnt-signalling has been linked to several pathological features of OA, cartilage degradation in particular (3). Wnts are a family of lipid modified and glycosylated ligands capable of activating several signalling cascades, upon interaction with the Frizzled (FZDs) family of receptors and a variety of co-receptors such as low-density Lipoprotein-related protein receptors (LRPs) and Receptor Tyrosine Kinase-like Orphan Receptors 1 and 2 (4). The most investigated branch of the Wnt-signalling in the context of cartilage biology and OA is the Wnt/β-catenin pathway. Overactivation of this pathway has been linked to cartilage degeneration in OA both in human (5) and in experimental animal models. Targeting of the Wnt signalling for therapeutic purposes has anyway remained a hurdle for the still incomplete understanding of the mechanisms regulating its activation as well as reciprocal modulation of β-catenin dependent and independent arms of the signalling (6).

We have recently discovered that Wnt3a can simultaneously mediate the activation of both the Wnt/β-catenin pathway and the Wnt/Calcium Calmodulin pathway in the articular chondrocytes (7). The two pathways are in a steady state equilibrium in physiological conditions and disruption of this homeostasis can lead to cartilage degeneration and OA development (7,8). Nonetheless, different transcriptional targets were downstream the activation of the two branches of the signalling (8).

Here, by further analysing our previous data, we discovered that microRNA-544a (miR-544a) is an important modulator of the Wnt-signalling in chondrocytes, whose activity is upregulated in the articular cartilage in OA, and targeting of this microRNA could represent a new strategy to halt tissue degeneration during the progression of the disease.

## Methods

### Human articular cartilage samples and chondrocytes isolation

Preserved (Mankin Score ≤4) and damaged (Mankin Score >4) human articular cartilage (AC) was collected from patients (male and female, age range 50-90) undergone total hip or knee replacement because of OA, following collection of informed consent. Surgeries were performed at the South-West London Elective Orthopaedic Centre. All procedures were approved by the London South East Research Ethics Committee (19/LO/0742). Chondrocytes were isolated as previously described (Nalesso et al., 2011;2021).

Samples used for the in-situ hybridisation experiment were isolated from human femoral cartilage obtained from study participants with knee OA undergoing total knee arthroplasty and recruited within the Longitudinal Evaluation in the Arthritis Program (LEAP) Addendum: Biomarker Exploration Analysis cohort (Division of Orthopaedics, Schroeder Arthritis Institute, University Health Network, Toronto). Surgeon-identified cartilages were collected from discarded tissues during total knee arthroplasty, under informed consent and under research ethics board approval (REB # 14-7592.22).

### Primary cells, cell lines, cell transfection and stimulation

Human articular chondrocytes (HACs) were cultured with complete media [(CM; DMEM/F-12, ThermoFisher Scientific, Waltham, MA, USA), supplemented with 10 % Fetal bovine serum (FBS; Gibco Invitrogen, Carlsbad, CA, USA)] and 1 % antibiotic antimycotic solution (AA, Sigma-Aldrich, St. Louis, MO, USA)) AT 37°C in a 5 % CO_2_ incubator (Binder, Tuttlingen, Germany). HACs at passages 0 to 2 were used for the further experiments.

C28/I2 human chondrocyte cell line were grown and expanded in CM. For experiments requiring differentiated cells, 80% confluent cells were cultured for 7 days in differentiating medium [D-MEM/F-12 1:1 plus GlutaMax, 1 % AA (Sigma-Aldrich), 1 % Insulin-transferrin-selenium liquid Media Supplement (ITS) (Sigma-Aldrich)].

HEK293FT human embryonal kidney cell line was grown and expanded in high glucose complete medium [(HG-CM; (DMEM High Glucose with L-Glutamine and Sodium Pyruvate, Biowest), supplemented with 1 % AA (v/v), 1 % MEM non-essential amino acids (v/v), and 10 % (v/v) FBS]. Experiments were conducted when cells reached 80 % confluence.

### Cell transfection

HACs or C28/I2 cells were transfected with hsa-miR-544a miRCURY LNA miRNA mimic (50nM; Qiagen) or negative control (NC) miRCURY LNA miRNA mimic (50nM; Qiagen) by using Lipofectamine 2000 (ThermoFisher Scientific) following manufacturer instructions for 4 hours at 37°C. After transfection, the cells were washed and incubated in CM with or without specific stimuli, as detailed in the “results” section. The sequence of NC has no homology to any known miRNA or mRNA sequences in murine or human. The sequences of miR544a mimic and of the NC can be found in Supplementary Table 1.

### In situ Hybridization (ISH)

Individual sections were deparaffinized and tissues were digested with Proteinase K (20 µg/ml; Qiagen) at 37°C for 20 min. Digoxigenin (DIG)-labelled probes were denatured at 90°C for 4 min and diluted with 1 x microRNA ISH buffer (Qiagen), following protocol by Endisha & Kapoor, 2021 (11). The sections were hybridized with hsa-miR-544a miRCURY LNA miRNA Detection Probe (150 nM; Qiagen), U6 snRNA (2 nM-positive control; Qiagen) or scramble control probe (40 nM – negative control; LNA-modified and 5’-and 3’-DIG-labeled antisense oligonucleotides) for 4 h at 55°C, using a humid StainTray. After hybridization step, stringent washes were performed with saline sodium citrate (SSC) 5x, SSC 1x and SSC 0.2x buffers. The sections were then treated with a blocking buffer (2 % sheep serum) for 15 min at room temperature (RT) and then incubated with anti-DIG-AP (Roche; 1:500) diluted in blocking buffer for 1 hour at RT. After that, the slides were incubated with AP substrate [4-nitroblue tetrazolium (NBT)-5-bromo-4-chloro-30-Indolyl-phosphate (BCIP) substrate; Vector Laboratories] in MilliQ water with 0.2 mM Levamisol for 3h at 30 ^O^C and then counterstained with Nuclear fast red (Sigma-Aldrich) for 3 min. Finally, sections were placed in water, dehydrated in alcohol, and mounted with VectaMount® Express Mounting Medium (Vector Laboratories). Images were acquired in a EVOS XL Core system microscope, using x10 and x40 objective lenses.

### Micromass cultures and Alcian Blue staining

Micromass cultures (MM) (25 x 10^4^ cells/well) were prepared as previously described (9). The MM were stimulated with recombinant human Wnt3a (100 ng/mL; R&D), or with recombinant human IL-1β (20 ng/mL; BioLegend) for 72h. Then, the MM were fixed and stained overnight with Alcian blue 8GS (Carl Roth, Karlsruhe, Germany) as previously described (12). Proteoglycans were extracted overnight with 6M guanidine hydrochloride (Sigma-Aldrich) and absorbance was read at 630 nm with a CLARIOstar spectrophotomer (BMG LABTECH, Offenburg, Germany). Absorbance values were normalized to protein content, determined by the bicinchoninic acid (BCA) assay, according to manufacturer instructions (ThermoFisher Scientific). Images of the micromasses were acquired at room temperature with a stereomicroscope (SZTL 350 Stereo Binocular Microscope, VWR®, Radnor, PA, USA).

### Immunocytochemistry

HACs (4 x 10^4^ cells/well) were seeded on 8-well chamber slides (Lab-Tek) and transfected with hsa-miR-544a mimic or NC (50 nM), as described above, and then stimulated for 24h with recombinant Wnt3a (100 ng/mL), KN92 (10 µM; EMD), KN93 (10 µM; EMD), or XAV-939 (10 µM), alone or in combination, as described in the individual experiments. After being washed in PBS, the cells were fixed in 4 % buffered PFA. Autofluorescence was quenched by washing the slides in 50 mM NH4Cl to minimise cell-specific autofluorescence and then blocked for 1 hour in blocking buffer (goat serum diluted 1:50 in PBS/0.5 % BSA). After blocking, samples were incubated overnight at 4°C with the following antibodies: anti–phospho-CaMKII rabbit polyclonal IgG antibody (Cell Signalling Technology), anti-beta-catenin (Cell Signalling Technology), anti-MMP-13 mouse monoclonal antibody (Santa Cruz Biotechnology) or anti-Aggrecan Neoepitope polyclonal antibody (Invitrogen), all diluted 1:100 in blocking buffer. Upon washing 3x in PBS, samples were incubated with a secondary Alexa Fluor 647-conjugated goat anti-mouse antibody (1:200; Invitrogen) for 1 hour at RT and washed 3x in PBS. Finally, the slides were stained with DAPI (Sigma-Aldrich). The slides were then mounted in Mowiol (Sigma-Aldrich) and images were acquired with a confocal fluorescence microscope (Nikon), using a 40X objective lens.

### Gene expression analysis

Total RNA (350 ng/sample) was extracted from HACs using TRIzol reagent (Invitrogen), according to manufacturer’s instructions. Samples were reverse transcribed to cDNA using a High-Capacity cDNA Reverse transcription kit (Applied Biosystems, Waltham, MA, USA). Quantitative PCR was performed using hot-start DNA polymerase (Qiagen, Hilden, Germany). The complete list of primers is presented at the Supplementary Table 2 (all primers were purchased from Sigma-Aldrich). Results were normalised for β-actin values. All reactions were performed in the CFX384TM Optics Module PCR machine (BioRad, Hercules, CA, USA).

To analyse miR-544a expression levels, samples were reverse transcribed using the cDNA miScript PCR Reverse Transcription Kit (Qiagen). The expression levels of precursor and mature miR-544a were evaluated by qPCR using miScript SYBR Green PCR Kit (Qiagen) and normalized for the housekeeping gene U6B small nuclear RNA (RNU6B). The primers were purchased from Qiagen (Catalogue number: MS00010087 for miR-544a; Catalogue number: MS00033740 for RNU6B) and the qPCR reactions were performed in the CFX96TM PCR machine (BioRad).

### Dual Luciferase reporter assay

Potential targets of miR-544a were predicted by using TargetScan release 7.2. Selection of targets genes was then performed through pathway analysis done with ConsensusPath database(13). Predicted pairing positions between miR-544a and 3’UTR of the selected genes were obtained using Target Scan and Microrna.org. 3’UTR regions of the transcripts for all the selected genes were downloaded in FASTA format from the Ensemble Genome Browser. Primers were specifically designed by using open-source Primer3Plus software to amplify a 0.5-1 kb region of 3’UTR containing the miR-544a binding sites. Extensions of 15 bases comprising the restriction site of the *Sac*I restriction enzyme (5’GAGCTC3’) were added to the 5’ end of the primer to obtain PCR amplicons homologous to 15 bases at each end of the pmirGLO Dual-Luciferase miRNA Target Expression Vector (Promega, Madison, WI, USA). Primer sequences are listed in Supplementary Table 3. Genomic DNA extracted from human chondrosarcoma SW1353 cells was used for the amplification of 3’UTR regions of the genes of interest, followed by the construction of the insertion in the vector and cell transformation.

After the validation of the selected putative targets of miR-544a, C28/I2 cells were co-transfected with miR-544a mimic (100 nM) or NC (100 nM) with the constructed pmirGLO Dual-Luciferase miRNA Target Expression Vectors (500 ng; Promega). After 48 hours, the media was discarded, and the reporter assay was performed by using the Dual Luciferase Reporter Assay System kit (Promega). Luminescence was measured at 560 nm with a CLARIOstar microplate reader (BMG LABTECH) and the results are expressed as mean ± SEM of Relative Luminescence units of Firefly luciferase normalized to *Renilla* luciferase of three independent experiments.

### TOPFlash reporter assay

C28/I2 and HEK293FT cells were co-transfected with SUPER8XTOP/FOPFlash TCF/LEF-firefly luciferase reporter vector (450 ng/well, Promega) and with the control vector expressing *Renilla* luciferase (50 ng/well) by using Lipofectamine 2000 (ThermoFisher Scientific), following manufacturer’s instructions. After transfection, the media was replaced, and cells were stimulated with recombinant Wnt3a (100 ng/mL; R&D Systems) or vehicle control for 48 hours. Luciferase activity was quantified using the Dual Luciferase Reporter assay system (Promega). Firefly luciferase activity was normalized by the *Renilla* luciferase activity. Results are expressed as fold increase of relative luminescence units in comparison to NC of three independent experiments.

### Statistical analysis

Statistical significance between two groups was assessed by unpaired two-tailed t-test and non-parametric data between two groups were analyzed by a Mann-Whitney post-test. For multiple comparisons, statistical analysis was performed by one analysis of variance (ANOVA), followed by Tukey post-test. A value of P ≤ 0.05 was considered statistically significant for all comparison tests. All the data in the graphs are expressed as mean ± SEM.

## Results

### Wnt3a promotes the expression of miR-544a in human articular chondrocytes

Our previous work showed that Wnt3a, a Wnt ligand whose expression is modulated in OA (7) can simultaneously activate both β-catenin dependent and independent branches of the Wnt signalling system in articular chondrocytes (7). To understand the specific transcriptional targets downstream the activation of the individual pathways, we performed a microarray analysis on human articular chondrocytes (HACs) stimulated with Wnt3a with or without the CaMKII inhibitor KN93 (8). Our analysis showed that the gene mostly upregulated in response to Wnt3a stimulation in a CaMKII independent manner was Axin2, a very well characterised modulator of the Wnt-β-catenin dependent pathway(14,15). The most downregulated gene was instead the precursor of miR-544a, a microRNA which had been shown to activate the Wnt-β catenin pathway in cancer (16–21), but whose role has not been previously characterised in the articular cartilage (Figure 1a). These data were further validated by qPCR (Figure 1b). We then tested whether Wnt3a stimulation of HACs could affect the expression of the mature form of the microRNA. Wnt3a stimulation promoted the upregulation of the mature form of miR-544a (Figure 1c). This might suggest that the biogenesis of this microRNA could be either regulated through a negative feedback loop mechanism or that the pre-miRNA degradation occurs faster than the degradation of the mature form, as previously described in a different biological context (22).

**Figure 1:**
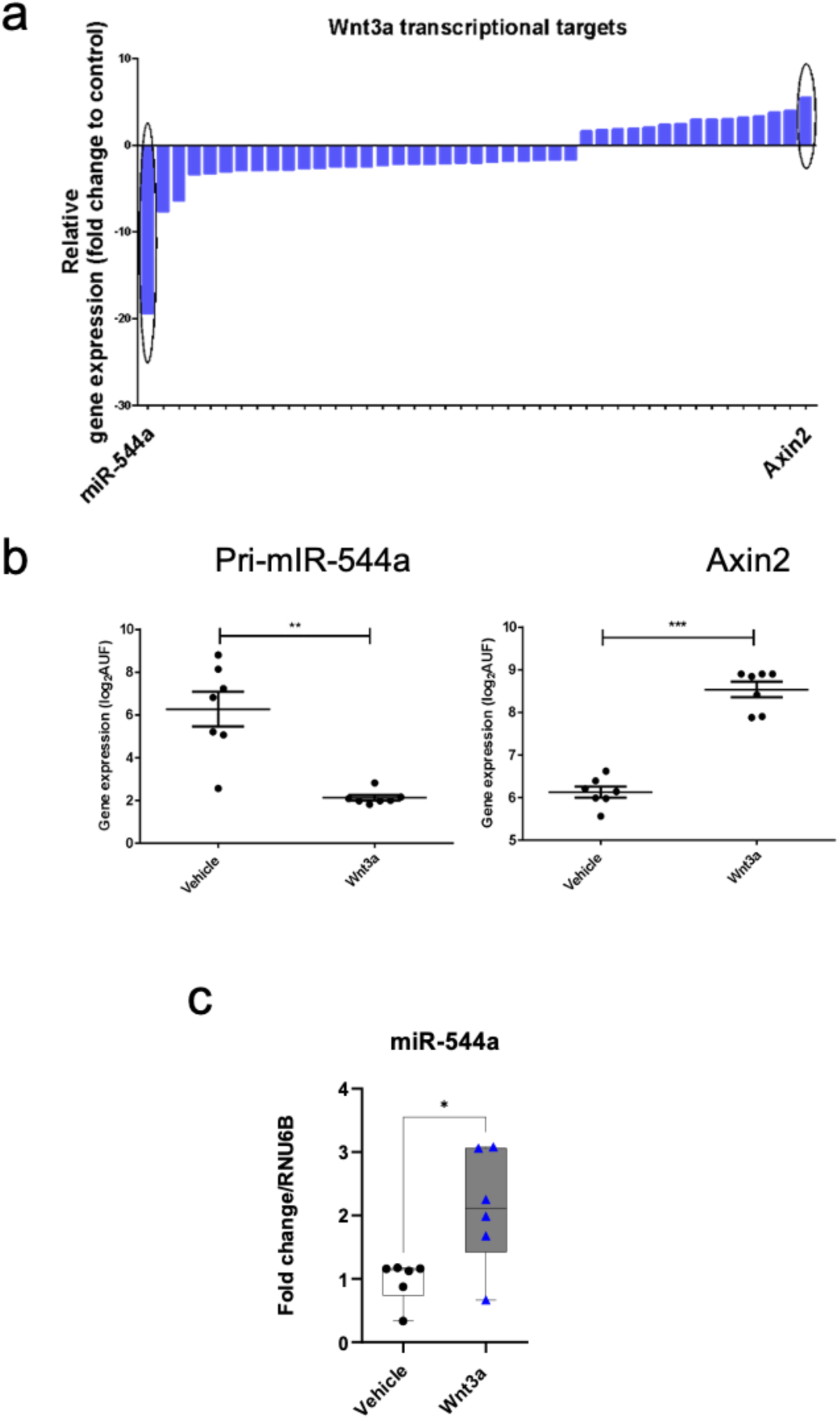
miR-544a is a transcriptional target of the Wnt-β-catenin signalling. (a) β-catenin dependent transcriptional targets of Wnt3a measured my microarray analysis (8). (b) mRNA expression levels of pri-miR-544a and Axin-2 in HACs stimulated with recombinant human Wnt3a (100ng/mL) for 24 hours, assessed by qPCR (n=6). (c) Expression levels of the mature form of miR-544a in HACs stimulated with 100ng/ml of recombinant human Wnt3a for 24 hours, assessed by qPCR (n=6). qPCR results for Axin-2 were normalised to the housekeeping gene β-actin and expressed as fold change in comparison to culture media or vehicle. qPCR results for miR-544a were normalised for the housekeeping gene RNU6B and expressed as fold change in comparison to vehicle. The fold change was calculated with the comparative 2^-ΔΔCt^ method. Data are shown as mean ± SEM. Statistical analysis was performed by unpaired two-tailed t-test followed by Mann-Whitney post-test. *p<0.05; **p<0.01; ***p<0.001.

### miR-544a modulates the activation of both β-catenin and CaMKII-dependent branches of the Wnt-signalling network

To define the molecular targets of miR-544a in articular chondrocytes, we performed a TargetScan analysis. Upon cross-referencing with previous literature, we focussed our attention on the 3p arm of miR-544a, as previous studies suggest this as an active form of this microRNA (23). The TargetScan analysis returned over 5154 putative target genes for miR-544a-3p. We then performed an over-representation analysis with ConsensusPath database software to understand the KEGG-annotated pathways potentially modulated by miR-544a. The analysis returned 69 pathways. Confirming previous literature, the Wnt pathway was among the pathways modulated by miR-544a. Putative targets of the pathway are listed in Figure 2a. Both components of the Wnt-β-catenin dependent and CaMKII-dependent branches of the Wnt signalling are included within this list. To validate the modulation of some of the components of the Wnt-β-catenin pathway in articular chondrocytes (Figure 2b), we cloned the 3’-UTR of Axin2, Cadherin 1 (CDH1), β-catenin, catenin beta interacting protein 1 (CTNNBIP1) and Glycogen synthase kinase 3 beta (GSK3β) or a scrambly rearranged sequence of their nucleotides as control upstream the luciferase expressing gene in pMirGlo-vectors. These were then transfected in the chondrocyte cell line C28/I2 (Figure 2c). Stimulation of the cells with a miR-544a mimic significantly decreased the relative luminescence emitted by the cells transfected with the original 3’-UTR sequences but not the one emitted by cells transfected with the mutated forms (Figure 2c). Modulation of the target genes was confirmed at mRNA level in HACs in Figure 2d. Interestingly, while the transcription of GSK3β seems to be downregulated by miR-544a in the reporter assay, the actual mRNA expression was upregulated by the microRNA in primary cells. This suggests the presence of additional molecular mechanisms controlling the expression of this key component of the Wnt-signalling in articular chondrocytes at the transcriptional level.

**Figure 2:**
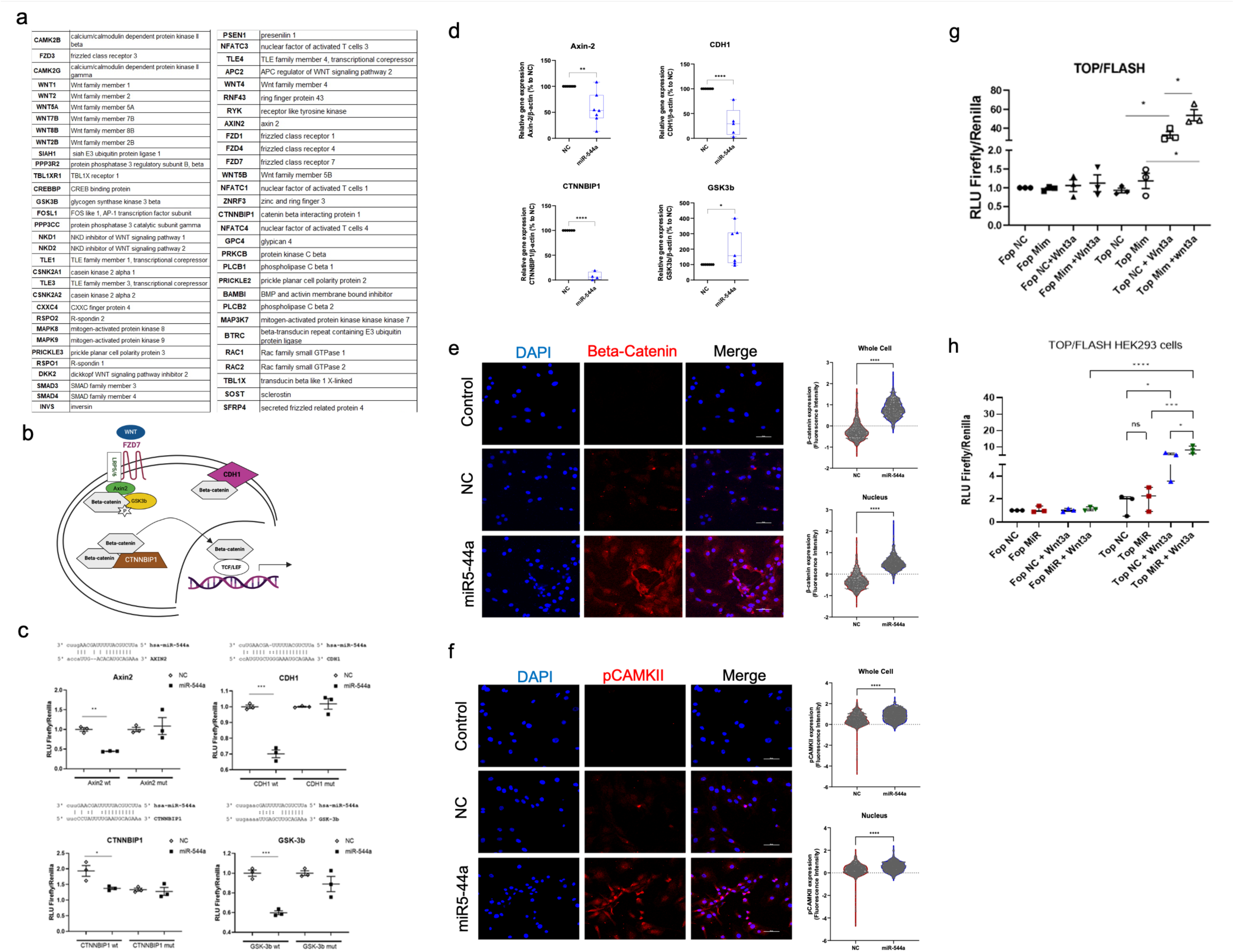
miR-544a modulates the activation of Wnt signalling pathways in articular chondrocytes. (a) Putative target genes of miR-544a associated with the activation of the Wnt pathway identified through TargetScan interrogation. b) Schematic representation of putative targets genes related to the β-catenin-dependent Wnt pathway. c) To validate some of these targets, 3’UTR-seed sequences of Axin-2, CDH1, CTNNBP1 and GSK3β or mutated sequences as control, were cloned in pmirGLO vectors upstream the Firefly luciferase gene. Confirming binding of miR-544a to these genes, reduced luminescence was recorded in C28/I2 cells transfected with miR-544a mimic (100 nM) and the vectors expressing the seed sequences 48h after transfection. No reduction in luminescence was reported in cells transfected with vectors expressing the mutated forms of the seed sequences. Results are expressed as mean ± SEM of Relative Luminescence units of Firefly luciferase normalized to Renilla luciferase (n=3). (d) mRNA expression levels of Axin-2, CDH1, CTNNBIP1 and GSK-3β in HACs transfected with miR-544a mimic (50nM) or negative control (NC, 50nM) for 3 hours, assessed by qPCR after 24 hours (n=6). qPCR results were normalised to the housekeeping gene β-actin and expressed relative gene expression (%) in comparison to NC. (d) Immunofluorescence analysis showing upregulation of β-catenin at protein level in HACs transfected with miR-544a mimic (50nM) in comparison to NC mimic (50nM). Fluorescence intensity of immunoreactive areas was quantified on the whole cell and nucleus after 24 hours upon transfection. (n=4). MiR-544a (50nM) enhanced the ability of Wnt3a (100ng/ml) to activate the SUPER8XTOPFlash reporter assay in (f) HEK293FT cells and (g) C28/I2 cells (n=3). (h) Immunofluorescence analysis showing upregulation of phospho-CamKII (pCamKII) expression in HACs response to transfection with miR-544a mimic in comparison to NC (both 50nM). Fluorescence intensity of immunoreactive areas was quantified on the whole cell and nucleus after 24 hours of transfection. (n=4). DAPI – positive staining for nuclei. Original magnification – x40, scale bar-50 µM. Data are shown as mean ± SEM. Statistical analysis was performed by one-way ANOVA or unpaired two-tailed t-test followed by Mann-Whitney post-test. *p<0.05; **p<0.01; ***p<0.001; ****p<0.0001.

To confirm the modulation of the Wnt-β-catenin pathway in response to miR-544a, we measured the expression of β-catenin in HACs stimulated with the miR-544a mimic. Indeed, β-catenin expression was upregulated by miR-544a and its transmigration to the nucleus was enhanced in comparison to cells stimulated with the negative control (Figure 2e). Furthermore, miR-544a enhanced the activation of a TCF/LEF reporter assay induced by recombinant Wnt3a, both in bovine primary chondrocytes and in HEK293 cells (Figure 2f and g). Finally, as some isoforms of CaMKII were also among the putative target genes for miR-544a, we tested whether stimulation of primary cells with miR-544a mimic could affect the activation of CaMKII. Phosphorylation of CaMKII was increased by stimulation of HACs with miR-544a, suggesting a role for miR-544a in the simultaneous modulation of separate branches of the Wnt-signalling (Figure 2e and h).

### miR-544a expression is upregulated in OA and by pro-catabolic cytokines

To investigate the biological role of miR-544a on cartilage homeostasis, we measured its expression levels in chondrocytes removed from preserved (Mankin score ≤4) and damaged (Mankin score >4) areas of articular cartilage removed from patients undergone total hip or knee replacement because of OA. miR-544a expression was significantly higher in cells isolated from damaged areas of the cartilage (Figure 3a). Interestingly, we noted that upregulation of the expression of the microRNA was not homogeneous within the tissue, but was higher in the deep layer, where the proteoglycan content was higher (Figure 3b). Interleukin-1 beta (IL-1β),-6 (IL-6) and transforming growth factor beta (TGFβ), pro-inflammatory and pro-fibrotic cytokines known to promote degeneration of joint tissues in the articular cartilage during the development of OA, increased the abundance of miR-544a in HACs (Figure 3c).

**Figure 3:**
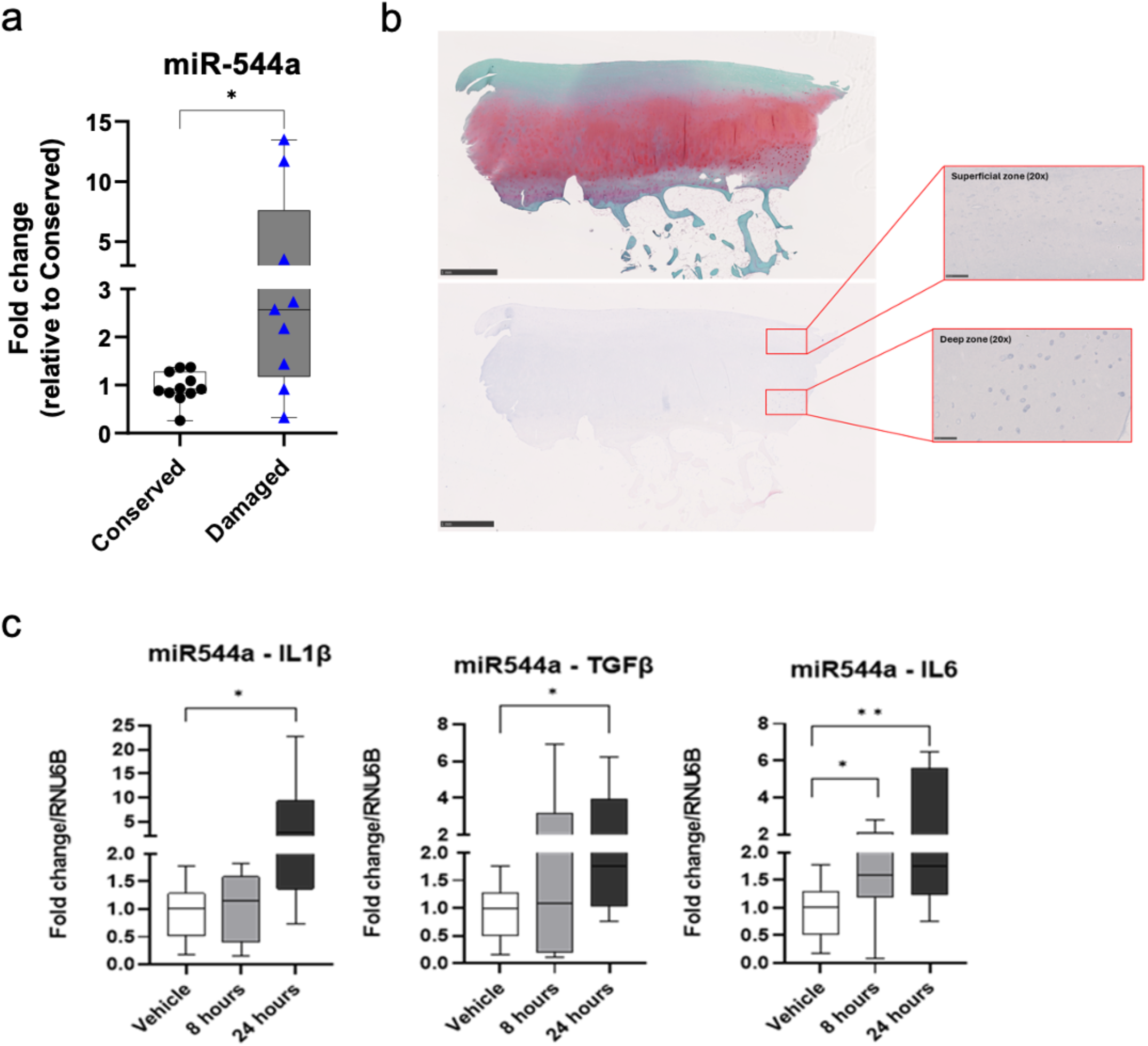
miR-544a is upregulated in OA and promotes catabolism in the articular cartilage. (a) Expression level of miR-544a in HACs isolated from conserved and damaged areas assessed by qPCR (n=4). (b) Representative image of in situ hybridisation for miR-544a in cartilage explants from the articular cartilage of patients undergoing joint replacement because of OA (n=6). (c) Expression level of miR-544a in HACs stimulated with IL-1β (20ng/mL), TGF-β (20ng/mL) or IL-6 (20ng/mL) for 8 hours or 24 hours, assessed by qPCR (n=4). qPCR results for miR-544a were normalised for the housekeeping gene RNU6B and expressed as fold change in comparison to vehicle. The fold change was calculated with the comparative 2^-ΔΔCt^ method. Data are shown as mean ± SEM. Statistical analysis was performed by one-way ANOVA followed by Tukey post-test or unpaired two-tailed t-test followed by Mann-Whitney post-test. *p<0.05; **p<0.01.

### miR-544a promotes catabolism in articular chondrocytes

We then investigated the effect of miR-544a on chondrocyte metabolism. Firstly, we measured proteoglycan content in micromass cultures of primary chondrocytes stimulated with Wnt3a in presence or absence of miR-544a. The microRNA promoted the downregulation of proteoglycans alone and enhanced the downregulation induced by the recombinant Wnt3a (Figure 4a and b).

**Figure 4:**
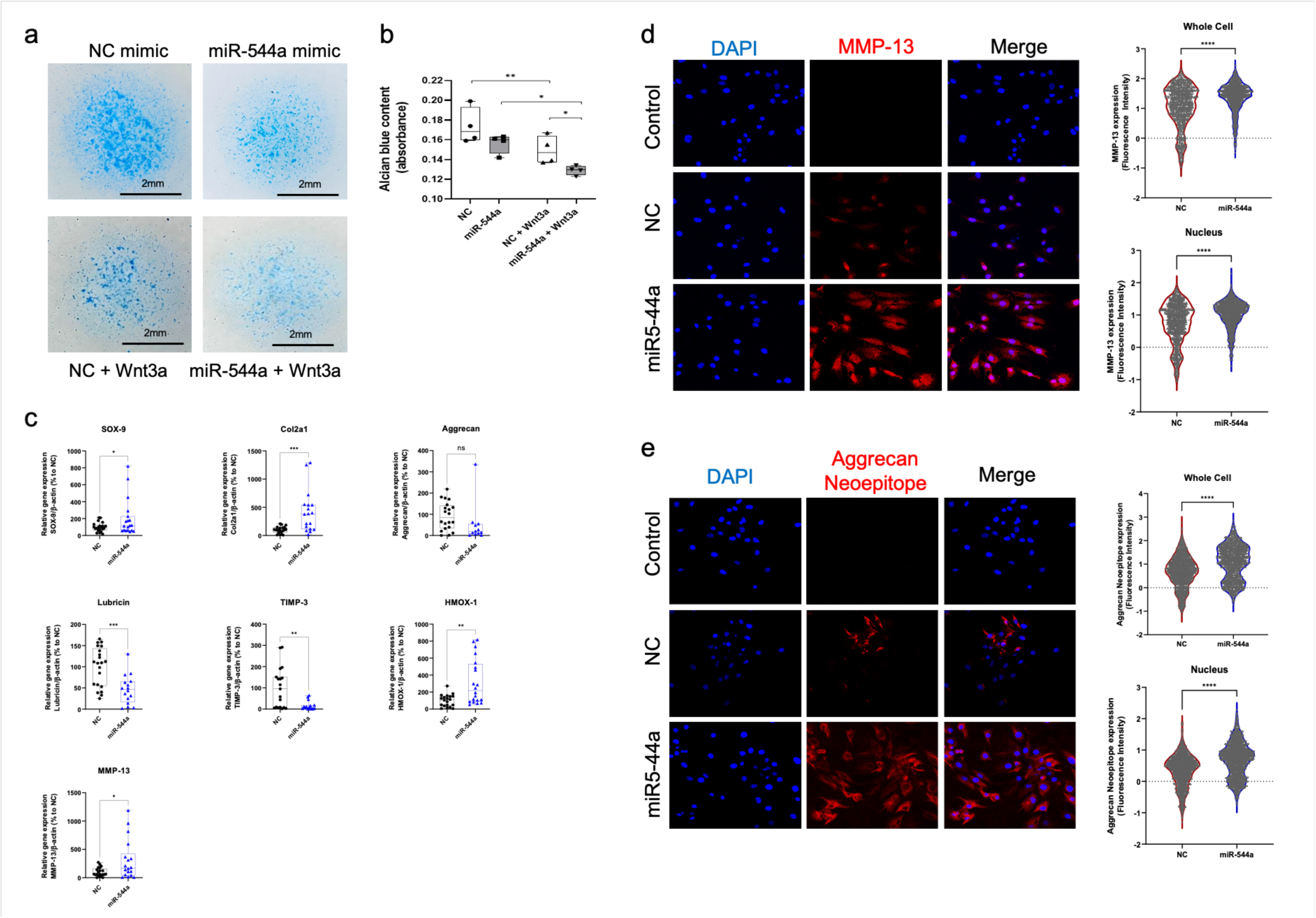
miR-544a modulates the expression of phenotypic genes and catabolic markers on HACs. (a, b) mir-544a mimic (50 nM), alone or co-stimulated with Wnt3a (100 ng/mL), reduced the accumulation of highly sulphated GAGs in chondrocyte micromasses, as assessed by Alcian Blue staining (n=4). (c) HACs were transfected with miR-544a mimic (50 nM) or NC (50 nM) in presence or absence of Wnt3a (100 ng/mL) for 24 hours and gene expression was assessed by qPCR (n=6). Individual gene expression levels were normalised by the expression levels of the house-keeping gene β-actin and expressed as the relative gene expression (% to NC). (d) MMP-13 and (e) Aggrecan neoepitope protein expressions were upregulated in HACs transfected with miR-544a mimic (50 nM) for 24 hours. Fluorescence intensity of immunoreactive areas was quantified on the whole cell and nucleus. (n=5). DAPI – positive staining for nuclei. Original magnification – x40, scale bar-50 µM. Data are shown as mean ± SEM. Statistical analysis was performed by one-way ANOVA followed by Tukey post-test or unpaired two-tailed t-test followed by Mann-Whitney post-test. *p<0.05; ***p<0.001; ****p<0.0001; ns=non-significant.

We then tested the effect of the microRNA on the expression of cartilage phenotypic markers and enzymes involved in the modulation of cartilage remodelling in OA. Stimulation of HACs with miR-544a mimic induced modulation of chondrocyte phenotypic markers, downregulation of the joint lubricant lubricin and of the metalloproteinase inhibitor Tissue Inhibitor of Metalloproteinase 3 (TIMP-3), whilst upregulating the expression of the hypertrophic marker MMP-13 (Figure 4c). miR-544a overexpression also promoted upregulation of Heme-Oxygenase 1 (HMOX1), a marker of cell oxidative stress (24) which we have previously shown to be a specific marker of the activation of the Wnt/CaMKII axis (8) (Figure 4c). We confirmed upregulation of MMP-13 at protein level by immunofluorescence (Figure 4d). Supporting a pro-catabolic role of miR-544a in the articular cartilage, we also showed that its overexpression can promote upregulation of Aggrecan neoepitope generated by Aggrecanase-mediated proteolytic cut (Figure 4e).

### miR-544a promotes catabolism through the activation of both the β-catenin and the CaMKII mediated branches of the Wnt signalling

As miR-544a could promote both the activation of the Wnt/β-catenin and of the Wnt/CaMKII mediated pathways, we investigated whether its catabolic activity was specifically mediated by one of the two branches. To this end, we stimulated HACs with the miR-544a mimic alone or in combination with inhibitors of the activation of CaMKII (KN93) or β-catenin (XAV939). Co-stimulation with both inhibitors rescued the upregulation of MMP-13 expression as well as the expression of Aggrecanase-cut aggrecan (Figure 5a and b). This seems to confirm an important modulatory role for miR-544a in the maintenance of the homeostatic balance between different branches of the Wnt signalling and that overactivation of both branches can lead to increased catabolism and pro-degenerative effects in the articular cartilage, as summarized in the schematic representation in Figure 5c.

**Figure 5:**
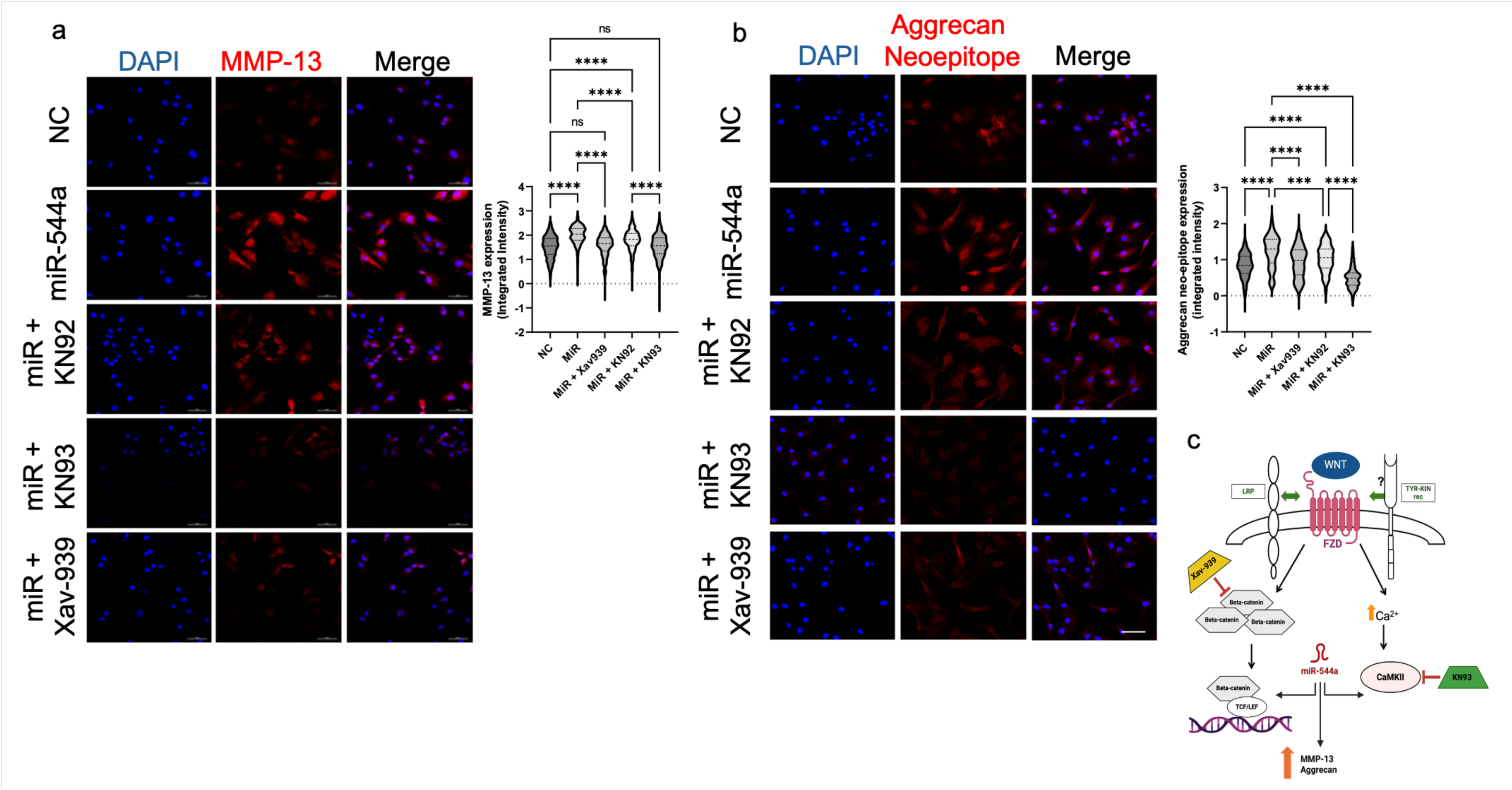
miR-544a promotes catabolism through activation of both β-catenin and CaMKII. (a, b) Immunofluorescence assay aimed to detect the expression of MMP-13 or MMP-cut aggrecan neoepitope in human articular chondrocytes (HACs) transfected with miR-544a mimic (50 nM) or NC (50 nM) in presence or absence of the β-catenin inhibitor Xav-939 (10 µM) or of the CaMKII inhibitor KN93 (10 µM) for 24 hours. Fluorescence intensity of immunoreactive areas was quantified on the whole cell (n=3). (c) Schematic representation of miR-544a modulation on the Wnt signalling pathway. DAPI – positive staining for nuclei. Original magnification – x40, scale bar-50 µM. Data are shown as mean ± SEM. Statistical analysis was performed by one-way ANOVA followed by Tukey post-test. ***p<0.001; ****p<0.0001; ns=non-significant.

## Discussion

Our previous work showed that Wnt ligands have the potential to simultaneously activate multiple signalling pathways with different biological outcomes. More specifically, we showed that Wnt3a could both activate the Wnt/β-catenin signalling pathway and the Wnt/CaMKII pathway driving both anabolic and catabolic changes in chondrocytes (7).

Here we investigated further microarray data previously generated, obtained by the stimulation of human articular chondrocytes with Wnt3a, to dissect specific transcriptional targets of the Wnt/β-catenin dependent pathway (9,10). Our data showed that the precursor of miR-544a was the transcriptional target mostly downregulated in a Wnt/β-catenin dependent manner by Wnt3a.

Interestingly here we showed that stimulation of chondrocytes also resulted in the upregulation of the mature form of miR-544a, suggesting that the biogenesis of this microRNA is regulated through a negative feedback loop mechanism or that the pre-miRNA degradation occurs faster than the degradation of the mature form, as previously described in a different biological context (22).

MiR-544a sits on the chromosome 14q32.31 region (Gene ID: 664613). Interestingly this chromosomic region has been associated with a susceptibility gene for OA (25) and cluster with other 19 micro-RNAs (mIR-134, -300, -323b, -376c, 376a-1 and 376a-2, -381, -382, -485, 487a and b, -539, -654, -655, -668, -889, -1185-1 and 2). Of these, 8 have already been associated with cartilage biology and/or the development of OA, albeit through interaction with different signalling cascades (26–33).

The role of this micro-RNA has mostly been described in cancer biology: experimental work pointed to miR-554a as a modulator of tumour growth and spreading of metastasis in different types of neoplasia (16–20,23,34,35).

The role of this microRNA in the articular cartilage and in OA has never been described before. Its modulation does not appear in previous transcriptomic studies aimed to identify biomarkers of disease. Nonetheless, we detected a higher abundance of the mature form of the microRNA in chondrocytes isolated from damaged areas of the articular cartilage of patients affected by OA. Within the damaged tissue, the expression of the microRNA localised in the areas where proteoglycan content was still abundant. These observations can lead to the speculation that the expression of this microRNA, and potentially many others, might be regulated in time and space during disease progression, and these factors should be taken more in consideration in the investigation of disease pathogenesis and to decrease data variability across donors.

Our findings confirm a role for miR-544a as an activator of the Wnt/β-catenin signalling. Yanaka and colleagues showed that miR-544a could induce the downregulation of CDH1 and Axin2, while favouring migration of β-catenin to the nucleus, and favouring epithelial-mesenchymal transition, thereby contributing to disease progression (21). Similarly, miR-544a was reported to downregulate the expression of GSK3β and promote the one of β-catenin sustaining the self-renewal ability of lung-cancer stem cells (36). Our data confirmed the important role of miR-544a in the modulation of the Wnt/β-catenin signalling in a non-cancer system, while adding complexity to its story, as we also demonstrated that this microRNA can also modulate the expression and the activity of components of the Wnt/CaMKII pathway in articular chondrocytes.

We have previously shown that the Wnt/β-catenin pathway and the Wnt/CaMKII pathways are in a steady-state equilibrium in the articular chondrocytes and their activation is reciprocally inhibitory (7).

We showed that a single Wnt-ligand could activate the two branches of the signalling in a concentration-dependent fashion. The data presented in this manuscript further highlight the strong connection between different arms of the signalling. As previously suggested by Amerongen and colleagues (37), we should start referring to the Wnt-signalling as a network and take a more holistic approach when taking in consideration its targeting for therapeutic purposes.

The importance and the complexity of investigating signalling pathways as part of cell networks rather than unidirectional mechanisms has been reviewed in a few previously published works. Jordan and colleagues suggested that signalling pathways share “junctions”, which act as integrators of signals, and “nodes”, in charge of splitting the signals and direct them to multiple outputs (38). While some of these key molecules have been identified, such as Protein Kinase A or Cdc42, the issue stands in how the signal within the network can be evaluated such as the appropriate response is then enabled. A physiological response is normally triggered when a signal comes through with the right intensity and for a specific but sustained duration. While this is easy to get through manipulation on in vitro conditions, it is not necessarily a reflection of real-life situations, where extracellular signals are mostly pulsatile and sub saturating in nature. In this context microRNAs such as miR-544a could really be crucial in stabilising signals and modulating their intensity to make them reach the right threshold, by acting on multiple components of the same network (38).

Integration of the Wnt signalling has been observed in different biological contexts, from development (39), where a concerted activation of different receptors and downstream targets lead to a precise spatiotemporal control of the neuroectoderm patterning, to cancer cells, where the Wnt/β-catenin dependent and the Wnt/Ca2+/CaMKII signalling have been proposed to be interdependent, as increased Ca^2+^ levels promoted translocation of β-catenin in the nucleus (40).

In conclusion, the results of this study add further layers of complexity in the modulation of the Wtn signalling in the articular cartilage, which however might help explaining the struggle in targeting this pathway for pharmacological purposes. A better understanding of the molecular switches responsible for the enhancement or lessening of the Wnt response in the cell could open new therapeutic opportunities for its pharmacological re-tuning to physiological level in pathological conditions such as OA.

## Funding

This work was supported by the Medical Research Council grant number MR/S008608/1, the Academy of Medical Sciences Springboard award SBF004/1112, the Longhurst Legacy fund, kindly donated by Sir Longhurst to the School of Veterinary Medicine to support teaching and research

## Notes

### Competing Interest Statement

The authors have declared no competing interest.

