## Supplementary Material for "microRNA-544a as a new modulator of the Wnt-signalling network in the articular cartilage and osteoarthritis"

**Supplementary material – Heluany CS et al 2024**

**Supplementary Table 1: Mimics sequences**

hsa-miR-544a miRCURY LNA miRNA Mimic

5'AUUCUGCAUUUUUAGCAAGUUC

Negative Control miRCURY LNA miRNA Mimic

microRNA strand: UCACCGGGUGUAAAUCAGCUUG

**Supplementary Table 2:** Primers sequences used for quantitative gene expression analysis.

| Gene | Sense Primer | Anti-sense Primer | Annealing Temperature |
|---|---|---|---|
| B-Actin | CACGGCTGCTTCCAGCTC | CACAGGACTCCATGCCCAG | 55°C |
| SOX-9 | GGCAAGCTCTGGAGACTTCTG | CCCGTTCTTCACCGACTTCC | 55°C |
| Aggrecan | GACTTCCGCTGGTCAGATGG | CGTTTGTAGGTGGTGGCTGTG | 55°C |
| Col2a1 | CTGCTCGTCGCCGCTGTCCTT | AAGGGTCCCAGGTTCTCCATC | 55°C |
| ADAMTS5 | GACCGATGGCACTGAATGTA | TGTACAGCTGGAGTTGTCTCCT | 55°C |
| MMP-13 | CCAGTTTGCAGAGCGCTACC | GACTGCATTTCTCGGAGCCT | 55°C |
| HMOX-1 | GAAAAGCACATCCAGGCAAT | ACTCAGGGCTTTTGGAGGTT | 55°C |
| TIMP-3 | GGAAGAGAGTACCGGCATCG | GATGTCCACTTGTGGGAGGG | 55°C |
| Lubricin | GGGAGATGTGGGGAAGGGTA | CCTTTACAGGAAAGCTCCGC | 55°C |
| Axin-2 | TACCGGAGGATGCTGAAGGC | CCACTGGCCGATTCTTCCTT | 60°C |
| CDH1 | ATACCCTTGTCCCCTGGGCTT | ACATTCGTCACTGCTACGTGT | 60°C |
| GSK3b | GGACTAAGGTCTTCCGACCC | TGTTAGTCGGGCAGTTGGTG | 55°C |
| CTNNBIP1 | GCGGTGACTCTCGGAGC | GCTGGGAGAAGTAGGAAGG | 60°C |

**Supplementary table 3:** Primers for 3'UTR sequences amplification

| *Gene* | Sense primer | Anti-sense primer | Length |
|---|---|---|---|
| ***Axin-2*** | 5’- **ctagttgtttaaacg** GCCTGTATTTGAGAGACTGCC-3’ | 5’- **gactcgaggctagcg** GGAGCCTCAGTCACAGGATC-3’ | 561 bp |
| ***CDH1*** | 5’- **ctagttgtttaaacg** CCATGTGCTGGGAAATGCAG-3’ | 5’- **gactcgaggctagcg** CCTAGTCAAGGTGGCCAGAC-3’ | 1151 bp |
| ***GSK-3β*** | 5’- **ctagttgtttaaacg** CTGCTGAGGTCAAAGGCTGA-3’ | 5’- **gactcgaggctagcg** TGCCAACAGACTCCACTTCC-3’ | 1022 bp |
| ***CTNNBIP1*** | 5’- **ctagttgtttaaacg** CTGCAAAGCCCTTGGAACAC-3’ | 5’- **gactcgaggctagcg** GTGAAGGCAGCAGCAAGTTC-3’ | 640 bp |
